# How drought and ploidy level shape gene expression and DNA methylation in *Phragmites australis*

**DOI:** 10.1101/2025.03.31.646440

**Authors:** Kristina Kuprina, Kerstin Haldan, Stepan Saenko, Mohamed Safwaan Gulam, Jürgen Kreyling, Martin Schnittler, Manuela Bog

## Abstract

Drought stress significantly affects plant physiology and growth, yet the molecular mechanisms underlying drought responses remain poorly understood. In this study, we investigate how tetraploid and octoploid *Phragmites australis* (common reed), a key species in wetland ecosystems and paludiculture, respond to drought at the transcriptional and epigenetic levels. Using RNA-seq, we identify changes in gene expression after 20 and 30 days of drought and assess methylation-sensitive amplification polymorphism (MSAP) over 50 days of drought. Transcriptomic analysis reveals that key drought-response genes are shared between ploidy levels, including those involved in the saccharopine pathway, water deprivation response, cell wall remodelling, and the mevalonate pathway. Drought supresses photosynthesis, with a pronounced down-regulation of the photosynthetic gene *PsbP*. Ploidy level influences gene expression under both drought and non-stress conditions, highlighting distinct adaptive strategies. In control samples, gene expression differed between ploidy levels, with octoploids up-regulating genes related to translation and metabolism, while tetraploids activate genes involved in cell wall modification and transmembrane transport. Prolonged drought increases DNA methylation variability, though no significant correlation is found between methylation levels and drought duration. Methylation differences are more pronounced between ploidy levels, with octoploids exhibiting lower overall methylation. These findings highlight the complex interactions between gene expression, epigenetic modifications, and polyploidy in drought response and provide a theoretical framework for future selection, hybridization, and conservation initiatives.

**Main Conclusion:** Key drought-response genes regulate saccharopine and mevalonate pathways, and cell wall remodelling. Ploidy level influences gene expression under drought and non-stress conditions. Octoploids overall exhibit lower methylation than tetraploids.

## Introduction

Drought is one of the most significant consequences of climate variability and anthropogenic effects (Haile *et al*. 2020). It is projected to increase in both frequency and intensity in the coming decades, particularly in mid-latitudes (Murray and Ebi 2012; Taylor *et al*. 2013; Rodell *et al*. 2018). These prolonged disruptions in water availability pose severe threats to ecosystems, particularly wetlands, which are highly dependent on stable hydrological conditions.

Among wetland plants, *Phragmites australis* (Cav.) Trin. ex Steud., or common reed, plays a significant role in wetland ecosystems, which is not only essential for maintaining biodiversity and ecosystem functions but also serves as a key species in paludiculture — a sustainable wetland use practice (Timmermann *et al*. 2006; Becker *et al*. 2020; Geurts *et al*. 2020; Lahtinen *et al*. 2022; Martens *et al*. 2023). This practice involves cultivating wetland plants such as *P. australis* under waterlogged conditions, helping to conserve carbon stocks, reduce greenhouse gas emissions, and support renewable biomass production, making it an effective tool for mitigating climate change (Wichtmann *et al*. 2020; Martens *et al*. 2022).

*Phragmites australis* stands out for its exceptional tolerance to environmental stress. Its adaptability is demonstrated by its broad ecological range, thriving in both freshwater and saline habitats. The species shows optimal salinity tolerance between 0 – 16 ppt, with certain lineages capable of enduring salinity levels up to 56 ppt (Lissner *et al*. 1999; Gao *et al*. 2012; Achenbach *et al*. 2013). Additionally, *P. australis* exhibits remarkable flexibility in water levels, with optimal growth occurring within a water table range of -30 cm to +70 cm and survival in water depths from -6 m to +2.3 m (Haslam 1972; Engloner 2009; Cui *et al*. 2010). Despite this adaptability, even brief periods of water deficiency can lead to reduced plant growth (Pagter *et al*. 2005). Rewetted peatlands, including those used in paludiculture, experience greater water table fluctuations (Kreyling *et al*. 2021). In combination with climate change, which lead to drought conditions in naturally moist habitats.

The high adaptability and phenotypic plasticity of *P. australis* is closely linked to its high genetic variation — traits commonly associated with widely distributed species (Clevering *et al*. 2001; Meyerson *et al*. 2016; Eller *et al*. 2017). Different genotypes of *P. australis* can significantly differ in their morphology and growth (Clevering 1999; Vretare *et al*. 2001; Haldan *et al*. 2023). These differences may persist even after several years of growth under similar conditions, with notable variation in biomass production, morphology, carbon/nitrogen dynamics, and ontogeny (Kühl *et al*. 1999; Pauca-Comanescu *et al*. 1999; Rolletschek *et al*. 1999; Koppitz *et al*. 2000; Hansen *et al*. 2007; Eller and Brix 2012; Haldan *et al*. 2023; Kuprina *et al*. 2023).

In addition to its high genetic variation, populations of *P. australis* comprise multiple phylogenetic lineages that have also independently developed different ploidy levels ranging from 3x to 12x, with tetraploid (4x) and octoploid (8x) cytotypes being the most prevalent (Clevering and Lissner 1999; Wang *et al*. 2024). Mixed cytotypes can also be found within a single population (Clevering and Lissner 1999; Lambertini *et al*. 2006; Hansen *et al*. 2007; Nakagava *et al*. 2013).

While genetic and ploidy variations play an important role in plant adaptation, gene expression regulation and epigenetic modifications are the major contributors to phenotypic plasticity in plants, allowing them to respond to diverse environmental conditions and stresses (Grativol *et al*. 2012). Drought-induced gene expression varies widely in plants, and the precise molecular regulation under water deficit conditions remains poorly understood, especially for non-model species (Reddi *et al*. 2004). It has been shown that *P. australis* can tolerate drought stress through osmotic adjustment and improved water-use efficiency, maintaining stable chlorophyll content and photosynthesis rates even under severe drought conditions (Pagter *et al*. 2005; Liu *et al*. 2018). However, no data exist on molecular mechanisms such as gene regulation and DNA modifications in *P. australis* under drought stress.

This study provides novel insights into the transcriptomic and epigenetic changes underlying drought responses in *P. australis*. Furthermore, in conjunction with the growth and physiology study by Haldan *et al*. (2025), it presents the first comparative analysis of the drought response of the two most common ploidy levels (tetraploid and octoploid) of *P. australis*. To account for potential variation among genotypes from different origins, we used regional pairs of tetraploid and octoploid genotypes collected from three distinct geographical regions in a single mesocosm experiment. After exposing plants to drought of varying durations, we analysed transcriptomic changes using RNA-seq and examined methylation patterns through methylation-sensitive amplification polymorphism (MSAP).

The aim of this study is to investigate the response of tetraploid and octoploid *P. australis* to drought stress by addressing the following questions: (i) Which genes and pathways are involved in the drought stress response? (ii) How does the drought influence methylation patterns in *P. australis*? (iii) Does ploidy level difference gene expression under non-stressed condition? (iv) Is there a difference in drought response between two ploidy levels?

## 2. Materials and Methods

### 2.1. Sample collection and drought experiment

The rhizome material was obtained from the live outdoor collection of *Phragmites australis* in the Aarhus University (Denmark, 56.228758 N, 10.127048 E) in November 2019. We selected six genotypes which were growing in the common garden under similar conditions for more than ten years and comprised three pairs of ploidy levels (tetraploid and octoploid) with different geographical origins: Lake Fertő in Hungary (Hu), Lake Razim in Romania (Ro) and Sakhalin Island in Russia (Ru) (Table 1; Lambertini *et al*. 2006). Ploidy level of the samples was proven by flow cytometry according to Kuprina *et al*. (2022).

**Table 1.** Samples of *Phragmites australis* used in this study*.

| Sample name | Days of drought | Ploidy | Genotype ID | Group: ploidy | Group: treatment | Number of raw read (million pairs) | Number of clean read (million pairs) | Mapping rate, % | Sequencing depth, x |
| --- | --- | --- | --- | --- | --- | --- | --- | --- | --- |
| c_Hu4x | 0 | tetraploid | Pa 664 HU | 4x | control | 59.06 | 39.32 | 75.69 | 62 |
| c_Hu8x | 0 | octoploid | Pa 663 HU | 8x |  | 52.59 | 40.18 | 74.64 | 55 |
| c_Ro4x | 0 | tetraploid | Pa 657 RO | 4x |  | 60.61 | 47.03 | 74.64 | 64 |
| c_Ro8x | 0 | octoploid | Pa 659 RO | 8x |  | 56.02 | 35.00 | 75.49 | 59 |
| c_Ru4x | 0 | tetraploid | Pa 205 RU | 4x |  | 40.85 | 26.61 | 74.98 | 43 |
| c_Ru8x | 0 | octoploid | Pa 178 RU | 8x |  | 56.38 | 43.83 | 74.10 | 59 |
| 2_Hu4x | 20 | tetraploid | Pa 664 HU | 4x | 20d | 14.79 | 9.39 | 72.63 | 16 |
| 2_Hu8x | 20 | octoploid | Pa 663 HU | 8x |  | 70.85 | 54.81 | 74.80 | 74 |
| 2_Ro4x | 20 | tetraploid | Pa 657 RO | 4x |  | 91.54 | 69.09 | 72.33 | 96 |
| 2_Ro8x | 20 | octoploid | Pa 659 RO | 8x |  | 40.73 | 29.45 | 68.11 | 43 |
| 2_Ru4x | 20 | tetraploid | Pa 205 RU | 4x |  | 56.61 | 44.63 | 73.18 | 59 |
| 2_Ru8x | 20 | octoploid | Pa 178 RU | 8x |  | 49.78 | 37.24 | 72.88 | 52 |
| 3_Hu4x | 30 | tetraploid | Pa 664 HU | 4x | 30d | 31.88 | 20.52 | 73.18 | 33 |
| 3_Hu8x | 30 | octoploid | Pa 663 HU | 8x |  | 16.63 | 11.01 | 74.10 | 17 |
| 3_Ro4x | 30 | tetraploid | Pa 657 RO | 4x |  | 116.42 | 110.08 | 73.74 | 122 |
| 3_Ro8x | 30 | octoploid | Pa 659 RO | 8x |  | 56.76 | 43.28 | 73.82 | 60 |
| 3_Ru4x | 30 | tetraploid | Pa 205 RU | 4x |  | 97.18 | 75.00 | 74.65 | 102 |
| 3_Ru8x | 30 | octoploid | Pa 178 RU | 8x |  | 68.34 | 52.63 | 73.01 | 72 |

\*Epigenetic study included additional samples of the same genotypes with 10, 40 and 50 days of drought

Meristematic propagation of rhizome buds was carried out at the Julius Kühn Institute in Braunschweig, Germany. The resulting plantlets were then transplanted to the Arboretum of Greifswald University (Germany, 54.092063 N, 13.410133 E) in June 2020. Subsequently, they were subjected to a mesocosm experiment from May to August 2021. Details about sample origin, experimental set up, growing conditions as well as measurements of the plants growth, morphology and photosynthetic rate can be found in Haldan *et al*. 2025. All the treatments ended on 27-08-2021. On this day, from 9:00 to 13:00, material for the transcriptome and epigenetic analysis was collected.

For transcriptome analysis, three levels of drought treatment were used: 0 days (control), 20 and 30 days of drought, giving 18 samples (Table 1). For each sample, tissue of three leaves was taken from each of three stems; a 0.5 cm^2^ piece of a second or third from the top fully developed and visually healthy leaf was cut in the middle position from the leaf base. The samples were incubated overnight at +6°C in RNAlater solution (Thermo Scientific, Waltham, MA, USA) and then stored at −20°C for one month.

For epigenetic analysis, five drought treatment levels were used: 0 days (control), 10, 20, 30, 40 and 50 days of drought yielding 30 samples. One leaf per plant was collected from three different stems and stored in a sealed plastic bag with silica gel for preservation.

### 2.2. RNA sequencing

Leaf material from one sample was pooled and homogenized in liquid nitrogen with mortar and pestle, then its total RNA was extracted using the innuPrep Plant RNA Kit (Analytik Jena, Jena, Germany) according to the manufacturer’s protocol. Directly after its extraction, RNA was examined for its concentration, purity and integrity using NanoDrop Lite spectrophotometer (Thermo Scientific) and a 1.5% (w/v) agarose gel. Then RNA was objected to DNase treatment using DNase I (Thermo Scientific) according to the manufacturer’s protocol and placed at -80°C. Samples were shipped on dry ice to Eurofins genomics (Konstanz, Germany) for the strand-specific cDNA library preparation and following sequencing on Illumina NovaSeq 6000 (Illumina Inc., San Diego, CA, USA) in S4 PE150 XP mode.

One RNA sample (c_Ro4x) was additionally sequenced using MinION Mk1C Model MIN-101C with Flow Cell (R10.4.1) and Direct RNA Sequencing Kit (SQK-RNA002) according to manufacturer’s protocol (all from Oxford Nanopore, Oxford, UK).

### 2.3. Transcriptomic data preprocessing

All bioinformatic steps were conducted using conda 23.9.0 and python 3.9.16.

Ribosomal RNA was filtered out from the raw Illumina reads using the program SortMeRNA 4.3.4 (Kopylova *et al*. 2012). Then the reads were trimmed for Illumina adapters and low-quality bases and the low-quality reads were discarded using trimmomatic 0.33 (Bolger *et al*. 2014) with the following parameters: *ILLUMINACLIP:TruSeq3-PE.fa:2:30:10, SLIDINGWINDOW:5:30, MINLEN:100.* Illumina and Nanopore Reads were taxonomically classified using the Kraken2 tool (Wood *et al*. 2019) and the KrakenTools 1.2 suite (Lu *et al*. 2022). Sequence databases used for classification included Standard (covering archaea, bacteria, viruses, plasmids, human, and vectors) and PlusPFP-16 (including protozoa and fungi), both released on 09-04-2024. Additional filtering of contamination was conducted by the NCBI Foreign Contamination Screen tool (Astashyn *et al*. 2024). Quality of the resulted reads was evaluated using programs FastQC 0.12.0 (Andrews 2010), MultiQC 1.14 and NanoPlot 1.43.0 (Ewels *et al*. 2016; De Coster and Rademakers 2023).

### 2.4. Assembly, annotation and quantification of transcripts

A flowchart of the data analysis workflow, scripts and MultiQC report for clean reads are available on GitHub: https://github.com/kuprinak/Phragmites_drought_RNAseq.

A hybrid *de novo* transcriptome assembly was performed in rnaSPAdes 3.15.4 (Prjibelski *et al*. 2020) using raw short reads of all control samples, as well as row long reads of the sample c_Ro4x. The resulting transcriptome was analyzed using rnaQUAST 2.2.3 (Bushmanova *et al*. 2016) to calculate basic statistics and predict gene numbers based on the GeneMarkS-T database. Additionally, the completeness of the transcriptome was assessed with BUSCO 5.4.4 (Manni et al. 2021) in the *euk_tran* mode, using the *poales_odb10* lineage dataset (creation date: 08-01-2024). Annotation of transcriptome was conducted using InterProScan 5.72-103.0 (Jones *et al*. 2014) against Pfam and SUPERFAMILY databases. Functional annotation has been done in KAAS 2.1 (Kyoto Encyclopedia of Genes and Genomes Automatic (KEGG) Annotation Server) (Moriya *et al*. 2007) using BLAST search against gene datasets of *Oryza sativa* subsp*. Japonica*, *Aegilops tauschii*, and *Zea mays*.

Quantification of reads was performed in Salmon 1.10.3 (Patro *et al*. 2017) via mapping clean reads against assembled transcriptome with the following parameters: -*l ISF, --gcBias, -- seqBias, --posBias, --validateMappings*.

### 2.5. Differential expression analysis

Analysis and visualization of differential expression of the transcripts was conducted in R 4.4.2 (R Core Team 2024) using RStudio 2023.12.1.402 (RStudio Team 2020). Results of the read quantification were imported to the R environment using *tximport* 1.32.0 package. The raw read counts were examined for variance distribution by plotting the variance against the mean count for each transcript. Raw read counts were then normalized using median of ratios method provided by the *DESeq2* 1.44.0 package (Love *et al*. 2014). To conduct a Principal Component Analysis (PCA), transformed read counts were additionally “regularized” via log transformation (function “rlog”), and then 300 transcripts with the highest variation were used for the analysis (function “plotPCA*”*). The matrix of these counts was used to draw a clustered heatmap in the package *pheatmap* 1.0.12 (Kolde 2018).

Transformed read counts were used for the calculation of Log2 fold changes (Log2FC) of the transcript abundances between groups of samples (Table 1) using the package *DESeq2*. To reduce the impact of random noise and control the false discovery rate, the resulted Log2FC were shrunk towards zero using method “ashr”. The transcripts with absolute Log2FC higher than 1.2 were assigned to differentially expressed genes (DEGs). The results were visualised using R packages *ggplot2 3.5.1*, *ggrepel* 0.9.6, and *ggvenn* 0.1.10 (Wickham 2016; Slowikowski 2024).

The enrichment analysis was conducted for DEGs grouped by assigned KEGG functions and Gene Ontology (GO) terms (biological processes, molecular functions and cellular components) using *clusterProfiler* 4.12.6 R package (Xu *et al*. 2024). The list of InterPro entries assigned to the GO terms was extracted from the Gene Ontology Resource webpage: https://current.geneontology.org/ontology/external2go/interpro2go (version date: 2024/04/08) (Mitchell *et al*. 2015). False discovery rate (FDR) adjustment was performed with p-value and q-value cutoffs set at 0.05.

### 2.6. Molecular analysis of MSAP

Various epigenetic mechanisms influence gene expression, but DNA methylation has been the primary focus of ecological studies (Schrey *et al*. 2013). Here, we employ a modified methylation-sensitive amplification polymorphism (MSAP) protocol to assess variation in DNA methylation at multiple sites of the genome (Reyna-López *et al*. 1997).

Total DNA was extracted from silica dry leaf material using the Mag-Bind Plant DNA DS Kit (OMEGA BioTek, Norcross, USA) according to the manufacturer’s protocol with the help of the KingFischer Flex Purification System (ThermoFisher, Waltham, USA). The concentration, purity and integrity of extracted DNA were measured by NanoDrop Lite spectrophotometer (Thermo Scientific, USA) and via electrophoresis on a 1.5% (w/v) agarose gel.

To control the error rate, 18% of the samples were processed in two technical replicates. Originally, more samples were included in the MSAP analysis. However, DNA extracts were processed in two batches, three months apart. Following data analysis revealed a strong batch effect, requiring the exclusion of one batch of samples.

DNA samples were digested separately using two restriction enzyme sets, each consisting of methylation-insensitive EcoRI and either methylation-sensitive MspI or HpaII. While EcoRI cuts its target site regardless methylation of cytosine, MspI does not cut when the inner cytosine is methylated, and HpaII does not cut when either or both cytosines are fully or hemi-methylated (see Roberts *et al*. 2007). Each 50 µL reaction contained 500 ng DNA, ddH₂O, 5 µL 10× Cut-Smart Buffer, 5 U EcoRI, and either 10 U MspI or 5 U HpaII (all from New England Biolabs (NEB), Ipswich, MA, USA). Reaction mixtures were incubated for 3 h at 37°C.

Resulted DNA fragments were ligated with the corresponding mix of adaptors. To prepare these mixes, two EcoRI adaptors (EcoRI-ad1: 5’-CTCGTAGACTGCGTACC-3’ and EcoRI-ad2: 5’-AATTGGTACGCAGTCTAC-3’, 100 µM each) or two HpaII/MspI adaptors (HpaII/MspI-ad1: 5’-CGATCAGGACTCATCG-3’ and HpaII/MspI-ad2: 5’-GACGATGAGTCCTGAT-3’, 100 µM each) were mixed 1:1 and incubated for 7 min at 95°C with the following 1 h cooling in a polystyrene box. Ligation mix of a total volume of 40 µl contained 20 µl restriction product, 4 µL 10x T4 ligation buffer (NEB), 3.3 U T4 ligase (NEB), 0.5 mM ATP (NEB), 40 pmol EcoRI-adaptors and 200 pmol HpaII/MspI-adaptors and ddH₂O. The mixtures were incubated for 8 h at 16°C. The ligation product was then diluted six-fold with ddH₂O.

Pre-selective amplification was conducted in two steps: First, 3.5 µL of diluted ligation product was mixed with 1 µL 10x PCR buffer, 0.88 µL 25mM MgCl_2_, 0.25 µL 10 mM dNTPs (all from Axon Labortechnik, Kaiserslautern, Germany), 3.15 µL ddH₂O, and 0.07 µL 5U/µL MolTaq DNA polymerase (Goffin Molecular Technologies, Beesd, Netherlands). The mix was incubated for 10 min at 60°C. After this step, we added 0.3 µL 10mM EcoRI-A primer (5’-FAM-GACTGCGTACCAATTC<u>A</u>-3’), 0.3 µL 10mM HpaII/MspI-A primer (5’-FAM-GATGAGTCCTGATCGG<u>A</u>-3’), 0.25 µL 10 mM dNTPs, 0.07 µL 5U/µL MolTaq DNA polymerase, and 0.07 µL ddH₂O. The amplification was conducted with SensoQuest thermocycler (Göttingen, Germany) with the following programme: (i) 2 min 30 s at 94°C; (ii) 10 cycles of amplification consisting of 30 s at 94°C, 30 s at 66 -56 ◦C (touch-down with -1°C per cycle) and min at 72°C; (iii) 20 cycles of amplification with 30 s at 94°C, 30 s at 56°C, and 2 min at 72°C; min final extension at 72°C. The resulting product was diluted 20-fold with ddH₂O.

Selective amplification was conducted with two sets of primers separately: (i) EcoRI-AGA with HpaII/MspI-ATG and (ii) EcoRI-ACT with HpaII/MspI-AGT. Primers had a FAM fluorochrome tag on the 5′ tail. Each 10 µL reaction mixture consisted of 0.9 µL of diluted pre-selective amplification product, 6.04 µL ddH_2_0, 1 µL 10x PCR buffer, 2.2 mM MgCl_2_, 0.5 mM dNTPs, 450 pmol EcoRI primer, and 160 pmol HpaII/MspI primer. Amplification conditions were the same as in the pre-selective amplification.

The samples were prepared for the capillary electrophoresis by adding 1.2 μl PCR product to 11.85 μl HiDi™ Formamide, 0.15 μl GeneScan™ 500 LIZ™ Size Standard and denaturation for 5 min at 95°C. Capillary electrophoresis was conducted with an ABI 3130XL Genetic Analyser. All from Applied Biosystems (Foster City, CA, USA).

### 2.7. Data analysis of MSAP

Detection and sizing of the peaks in resulted electropherograms were conducted via the program PeakScanner 2.0 (Applied Biosystems) with the following settings: full size range, minimum peak height of 50, light peak smoothing and “local Southern” size calling method. Output peak size table was further processed using R package *RawGeno* 2.0-2 (Arrigo *et al*. 2012) and R 3.6.2. Binning was conducted manually; peaks were scored in a range from 60 to 500 bp with the lowest fluorescence of 300 RFU. A combined 0/1 matrix of distance was created for both primer pairs and restriction enzymes.

The resulting matrix was analysed using the R package *msap* 1.1.8 (Pérez-Figueroa *et al*. 2013) in R 4.2.2. The package compared fragment presence between enzyme pairs (MspI vs. HpaII) to assess the methylation sensitivity of the corresponding restriction sites. It then estimated methylation levels across sample groups (ploidy levels and treatments), categorizing them as: (i) unmethylated, (ii) hemimethylated, (iii) methylated at the internal cytosine, or (iv) fully methylated (or absent target site). We tested Pearson’s correlation between drought duration and level of unmethylation or full methylation. Additionally, using the *msap* package, we performed molecular variance analysis (AMOVA) with 10,000 permutations and principal coordinate analysis (PCoA) based on Euclidean distances.

## 3. Results

### 3.1. RNA library

Illumina sequencing produced a total of 1,037.1 M raw read pairs with the average sequencing depth of 60x per sample (Table 1). After trimming and filtering out rRNA and contamination, 690.0 M clean read pairs were left (average ± SD: 38.3 ±18.5 M read pairs; Table 1). All sample libraries successfully passed the assessments of Phred read quality, per-base N content, adaptors and overrepresented sequences content.

Nanopore sequencing of the sample c_Ro4x generated 618.3 K reads with N50 length of 1.35 kb. After the removal of contaminating reads (mostly comprising DNA of Saccharomycetes), 273,336 reads were left (N50 read length = 918 bp, the longest read = 132 Kb).

### 3.2. Transcriptome

The assembled transcriptome comprised 258.76 Mb across 191,527 sequences, with an N50 length of 2,279 bp. BUSCO estimated the transcriptome completeness at 90.7%, while rnaQUAST predicted 88,348 genes. InterProScan successfully annotated 93,050 transcripts, with 92.5% assigned to 5,652 unique InterPro entries. Additionally, all transcripts were mapped to either one of 5,218 unique Pfam entries or 1,095 unique SUPERFAMILY entries. Functional annotation identified 21% of the transcripts as associated with 750 distinct KEGG identifiers.

The MultiQC reports of the quality of raw and clean reads can be found on GitHub: https://github.com/kuprinak/Phragmites_drought_RNAseq. Clean short and long reads, as well as assembled transcriptome, can be found on NCBI (PRJNA1195549).

### 3.3. Differential gene expression

Mapping of the clean reads on the transcriptome resulted on average of 73.66% of mapping rate (Table 1). PCA showed that the first component axis explains 47% of the variance and the second 11% (Fig. 1). Samples along the first axis were separated into two clusters: control and treated samples. Along the second axis, samples were separated by origin: from Europe (Hu and Ro) and East Asia (Ru).

**Figure 1.**
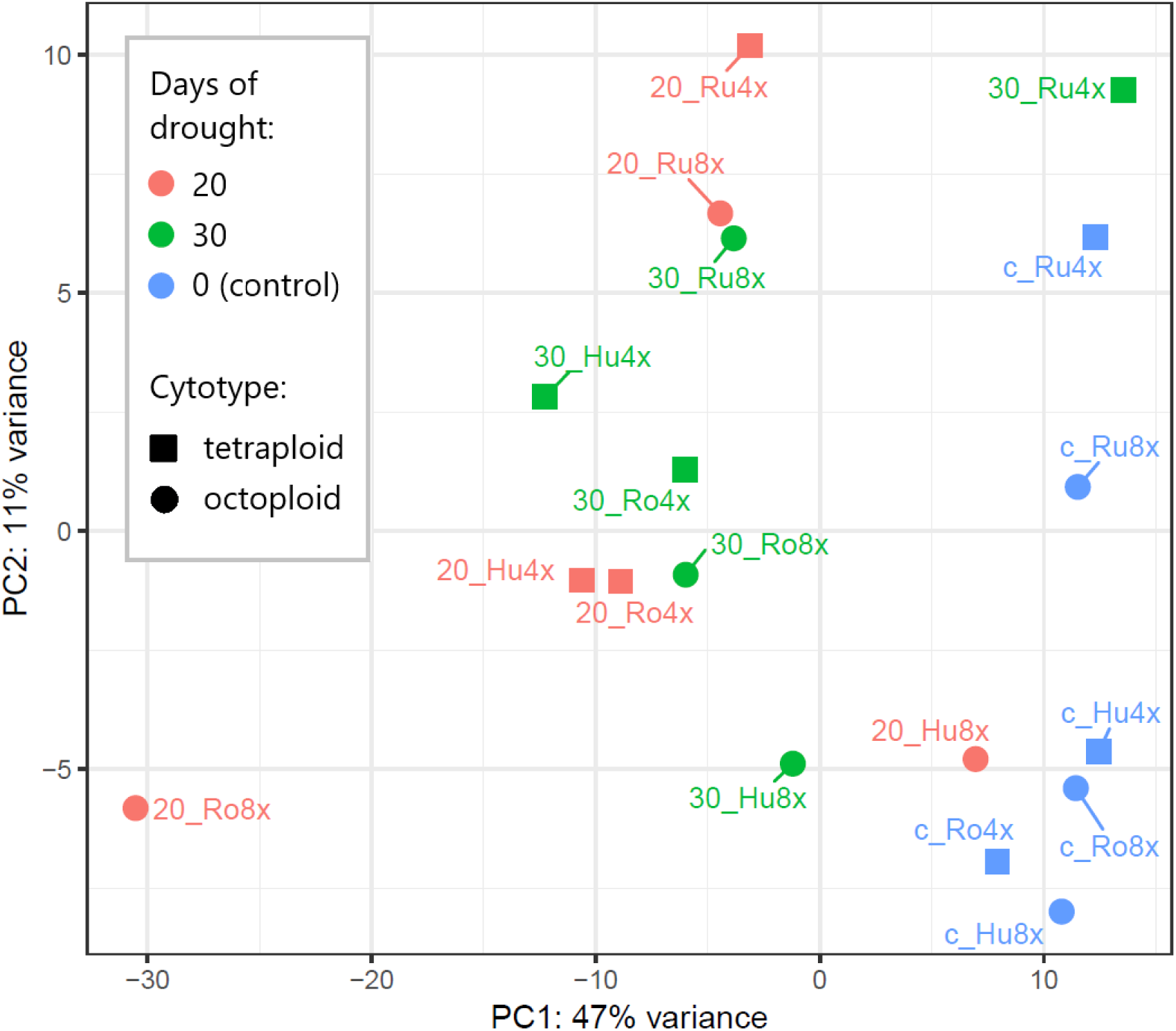
Ordination diagram of PCA of normalized counts of 300 most variable transcripts. Transcriptomic data were obtained for 18 leaf samples of *Phragmites australis* of two cytotypes (tetraploid and octoploid), three origins (Hu, Ro, Ru) which were subjected to drought treatments of 0, 20, or 30 days.

Differential gene expression analysis was conducted using 4,788 transcripts. The full list of Log2FC, p-values and adjusted p-values for each transcript and each comparison can be found in Suppl. Table 1. The number of DEGs varied across the comparisons of sample groups. After 20 days of drought treatment, 135 genes were up-regulated, and 43 were down-regulated compared to the control samples (Fig. 2a). By the 30th day of drought, 81 genes were up-regulated, and 126 were down-regulated (Fig. 2a). Between the 20- and 30-day drought treatments, 60 (39%) up-regulated and 27 (19%) down-regulated genes were shared (Figs. 2b, 2c). Only one down-regulated gene and no up-regulated genes were identified when comparing samples from 30 days to those from 20 days of drought.

**Figure 2.**
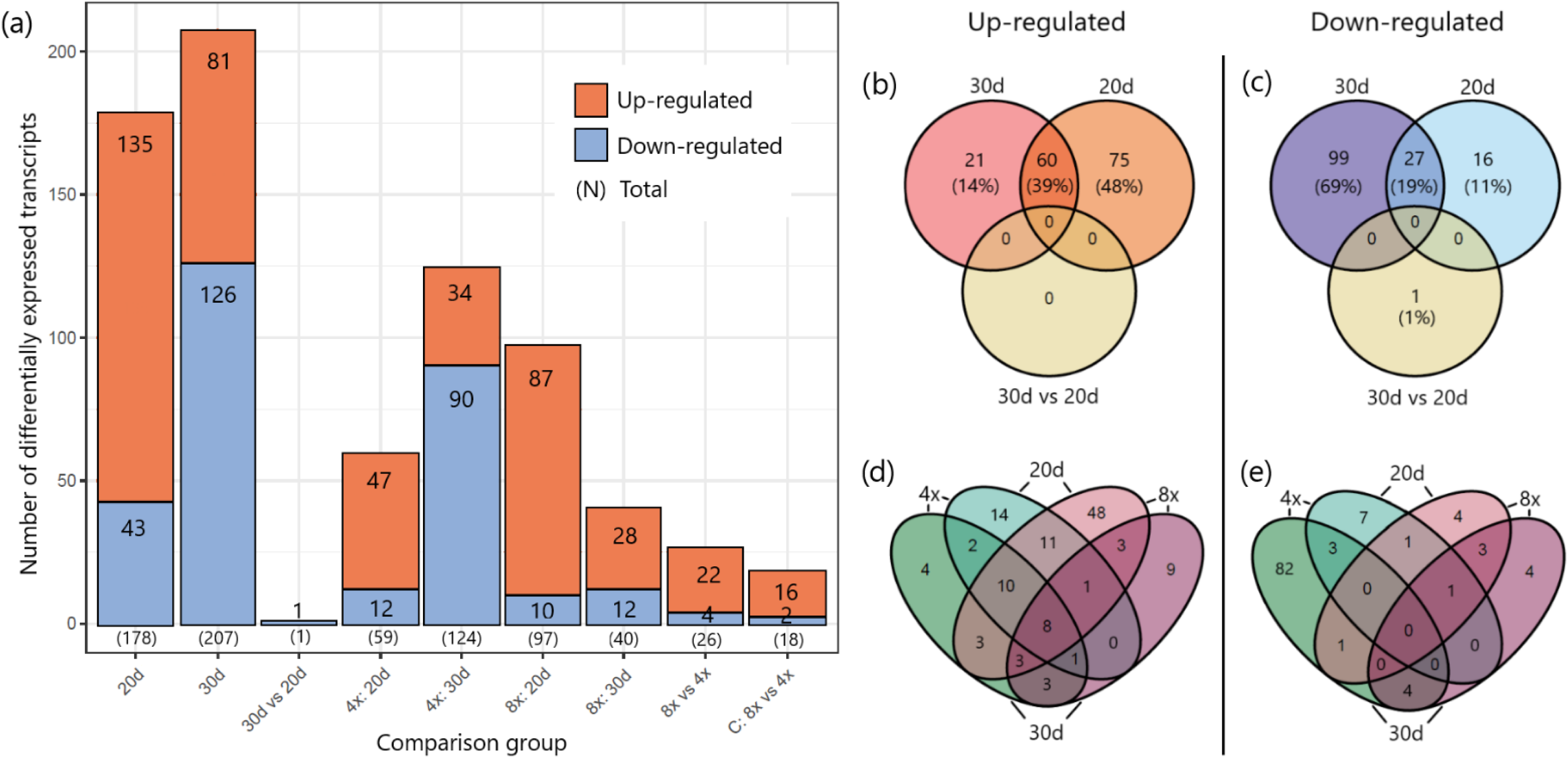
(a) Total numbers of differentially expressed genes across nine comparisons. (b-e) Venn diagrams displaying unique and shared (b, d) up-regulated and (c, e) down-regulated genes, with (b, c) showing all ploidy levels and (d, e) focusing on comparisons between tetraploid (4x) and octoploid (8x) samples separately. Transcriptomic analyses were conducted on 18 *Phragmites australis* leaf samples, comprising three tetraploid and three octoploid genotypes. Samples of each genotype was subjected to drought treatments of 0 days (control), 20 days (20d), and 30 days (30d). Differential gene expression analysis was performed on 4,788 transcripts.

Among all DEGs, eight genes were up-regulated in both drought treatments and both ploidy levels. They encoded: alpha-aminoadipic semialdehyde synthase (*AASS*; IPR007886), late embryogenesis abundant protein 1 (*LEA1*; IPR005513), dehydrin (IPR000167), saccharopine dehydrogenase (*SCCPHD*; IPR032095), heat shock associated protein 32 (*HSA32*; IPR003830), heparan-α-glucosaminide N-acetyltransferase (*HGSNAT;* IPR012429), glyoxalase (*GLO*; IPR029068), and domain of unknown function 599 (*DUF599*; IPR006747). Among these eight genes, seven showed Log2FC > 2 in at least one comparison (Fig. 3a). The BLASTN for three up-regulated genes with unknown function DUF599, DUF677, and DUF163 found the closest match with *P. australis* uncharacterised mRNA.

**Figure 3.**
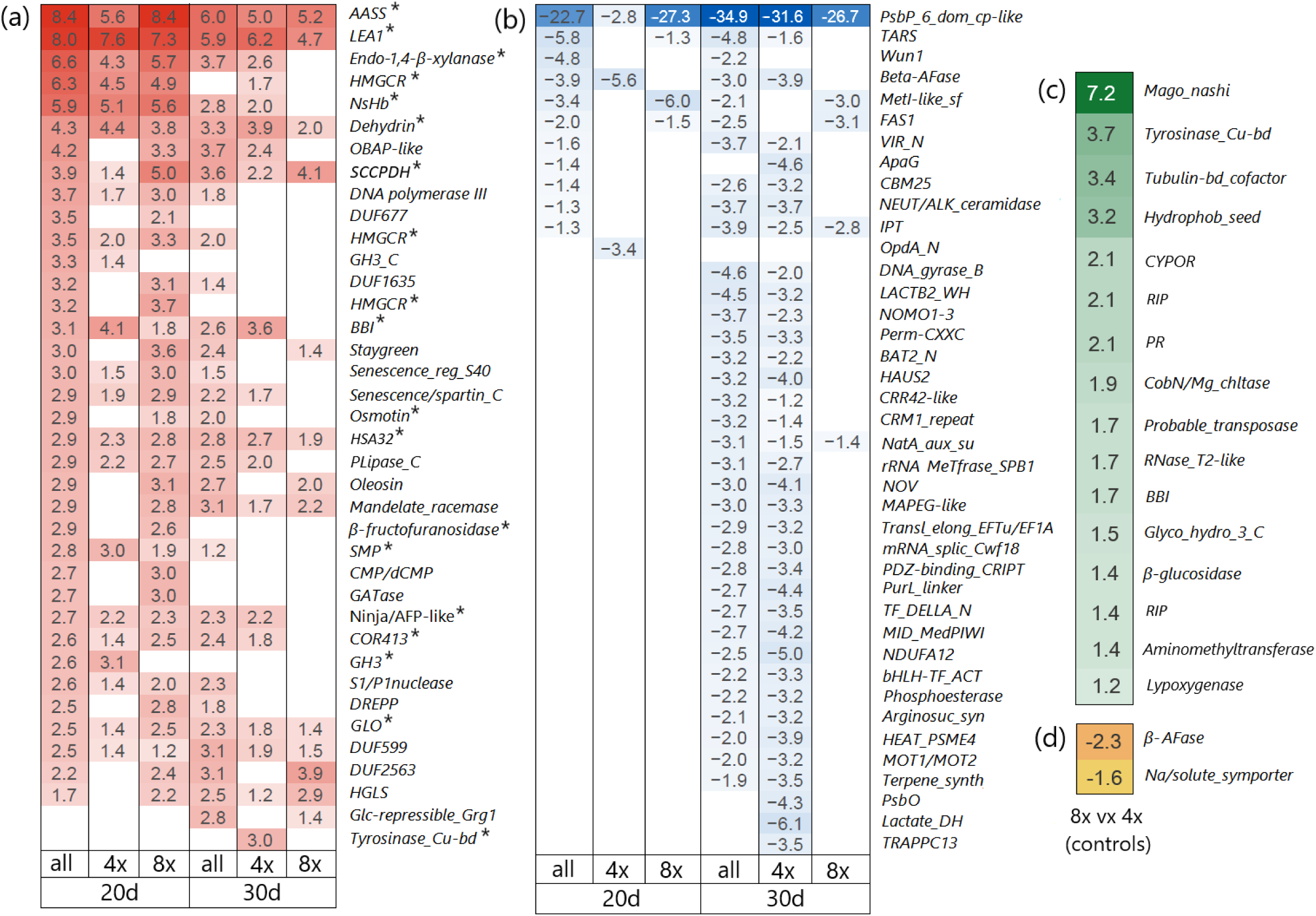
Heatmaps with the (a) up-regulated genes with Log2 fold change > 2.0 and with (b) down-regulated transcripts with Log2 fold change < -3.0 in *Phragmites australis* leaf samples after 20 days (n = 6) and 30 days (n = 6) of drought compared to corresponding control samples (zero days of drought, n = 6). Heatmaps (c) with all up-regulated and (d) all down-regulated transcripts identified in control octoploid samples (8x, n = 3) compared to control tetraploid samples (4x, n = 3). Numbers represent Log2 fold changes for genes with significant adjusted *p*-values (α = 0.05). Differential gene expression analyses were performed on 4,788 transcripts. *\** - genes putatively involved in the drought response

No genes were consistently down-regulated across both drought treatments and ploidy levels. The most highly down-regulated gene after both 20 and 30 days of drought encoded the domain of unknown function (*DUF3007* = IPR021562, up to 34.9-fold) (Fig. 3b). BLASTN search identified its closest match with the psbP domain-containing protein 6, chloroplastic-like (LOC133908951) of *Phragmites australis*.

A distinct pattern of transcripts expression appeared when comparing samples with different ploidy levels (Fig. 2a). Tetraploid samples exhibited more pronounced expression changes on the 30th day of drought, with 59 upregulated and 124 downregulated DEGs, whereas octoploid samples showed greater expression changes on the 20th day, with 97 upregulated and 40 downregulated DEGs. Tetraploid samples had the highest number of unique down-regulated transcripts (89), whereas octoploid samples exhibited the highest number of unique up-regulated transcripts (57) under drought stress (Figs. 2d, 2e).

The comparison of octoploid and tetraploid controls revealed 16 up-regulated transcripts and two down-regulated transcripts in octoploid samples (Fig. 2a; Figs. 3c, 3d). The most highly up-regulated genes (with the Log2FC > 2) encoded Mago Nashi protein (IPR004023, 7.2-fold), tyrosinase (Cu-binding domain, IPR002227, 3.7-fold), tubulin binding cofactor (IPR012945, 3.4-fold), hydrophobic seed protein domain (IPR027923, 3.2-fold), NADPH-cytochrome P450 oxidoreductase (CYPOR, IPR003097, 2.1-fold), ribosome-inactivating protein (RIP, IPR036041, 2.1-fold), and pathogenesis-related protein (PR, IPR014044, 2.1-fold). Two down-regulated genes encoded β-L-arabinofuranosidase (β-AFase, IPR012878, 2.3-fold) and sodium/solute symporter (IPR001734, 1.6-fold).

### 3.4. Enrichment analysis

Eleven GO entries were consistently enriched in the samples after 20 and 30 days of drought: eight were linked to up-regulated transcripts and three to down-regulated transcripts (Table 2). For KEGG enrichment analysis, three genes were up-regulated after both 20 and 30 days of drought: *cathepsin B* (K01363, involved in lysis), *acetyl-CoA synthetase* (K01895, associated with glycolysis/gluconeogenesis, pyruvate metabolism, glyoxylate and dicarboxylate metabolism, and propanoate metabolism), and *PPIP5K1* (K13024, involved in the phosphatidylinositol signalling system within environmental information processing). The full list of enriched GO and KEGG entries can be found in Supplementary File 1.

**Table 2.**
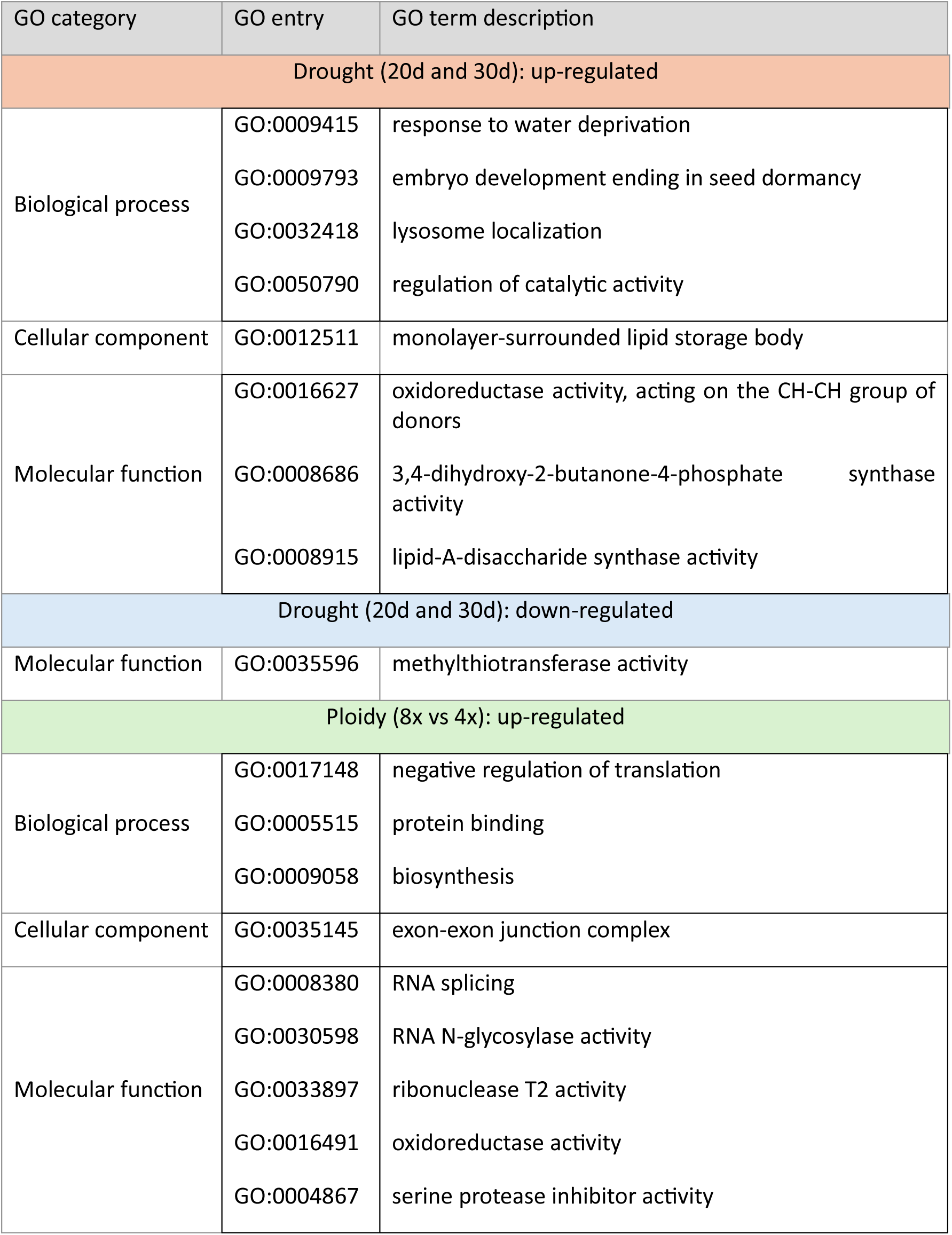
Enriched Gene Ontology (GO) entries identified in *Phragmites australis* leaf samples after 20 days (n = 6) and 30 days (n = 6) of drought compared to control samples (zero days of drought, n = 6), as well as in octoploid samples (n = 3) compared to tetraploid samples (n = 3).

| GO category | GO entry | GO term description |
| --- | --- | --- |
| Drought (20d and 30d): up-regulated |  |  |
| Biological process | GO:0009415 | response to water deprivation |
|  | GO:0009793 | embryo development ending in seed dormancy |
|  | GO:0032418 | lysosome localization |
|  | GO:0050790 | regulation of catalytic activity |
| Cellular component | GO:0012511 | monolayer-surrounded lipid storage body |
| Molecular function | GO:0016627 | oxidoreductase activity, acting on the CH-CH group of donors |
|  | GO:0008686 | 3,4-dihydroxy-2-butanone-4-phosphate synthase activity |
|  | GO:0008915 | lipid-A-disaccharide synthase activity |
| Drought (20d and 30d): down-regulated |  |  |
| Molecular function | GO:0035596 | methylthiotransferase activity |
| Ploidy (8x vs 4x): up-regulated |  |  |
| Biological process | GO:0017148 | negative regulation of translation |
|  | GO:0005515 | protein binding |
|  | GO:0009058 | biosynthesis |
| Cellular component | GO:0035145 | exon-exon junction complex |
| Molecular function | GO:0008380 | RNA splicing |
|  | GO:0030598 | RNA N-glycosylase activity |
|  | GO:0033897 | ribonuclease T2 activity |
|  | GO:0016491 | oxidoreductase activity |
|  | GO:0004867 | serine protease inhibitor activity |

The comparison of octoploid control samples with tetraploid control samples revealed nine up-regulated GO entries (Table 2) and one up-regulated KEGG entry, *lipoxygenase* (K00454), involved in the linoleic acid metabolism pathway.

### 4.5. MSAP

As a result, 32 samples in 252 loci were screened for methylation patterns. The detected error rate was 8.3% for HpaII and 7.0% for MspI in primer set 1, and 4.7% for HpaII and 6.7% for MspI in primer set 2.

No significant difference between drought treatments was found by AMOVA (df = 5, ΦST = 0.0027, *p*-value = 0.4330). Similarly, no significant correlation was found between drought duration and level of unmethylation (r = 0.67, *p*-value = 0.1451) or full methylation (r = -0.64, *p*-value = 0.1727). PCoA did not reveal separation of samples of any drought treatment but showed the increase in variation of methylation pattern with the increase of drought duration (Fig. 4a).

**Figure 4.**
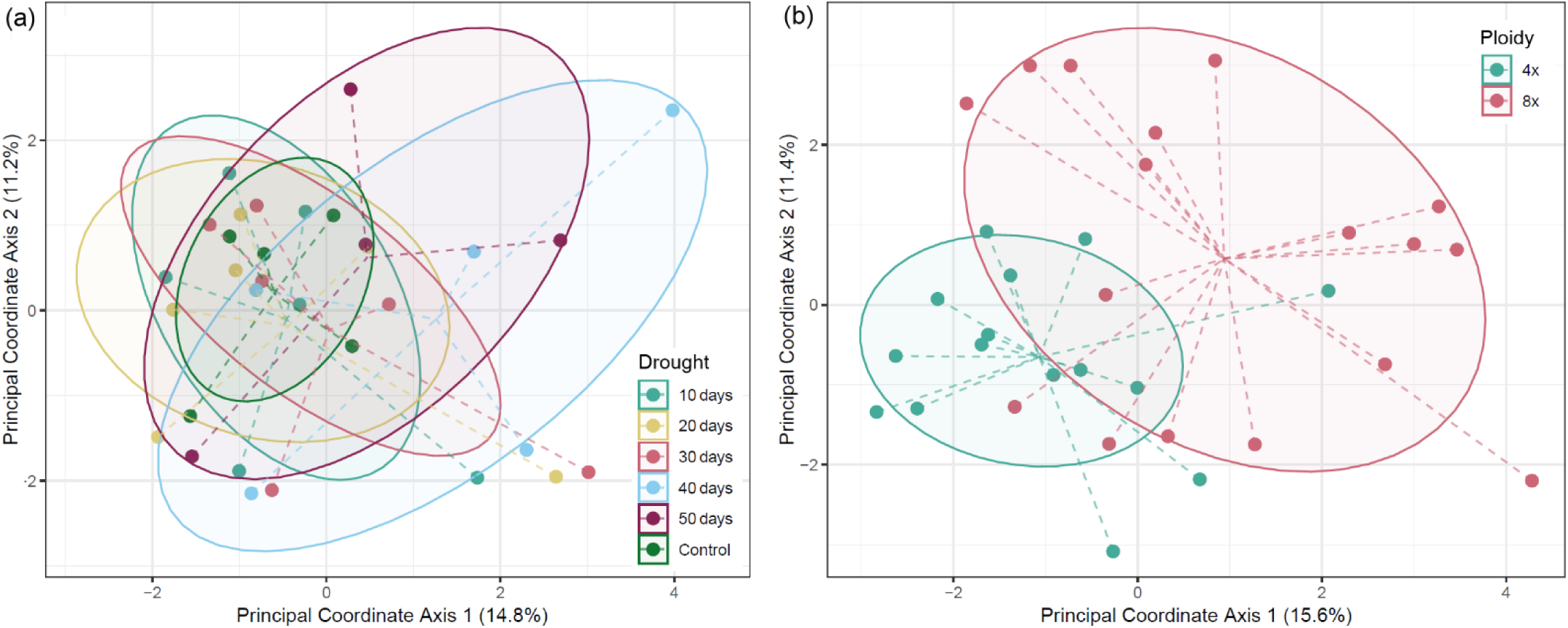
Ordination diagram of PCoA calculated on 248 methylation-sensitive loci for 32 leaf samples of *Phragmites australis* (a) under six different drought treatments and (b) with two ploidy levels. Ellipses depict one standard deviation of the data (α = 0.68).

AMOVA revealed significant differentiation between tetraploid and octoploid samples (df = 1, ΦST = 0.1153, p = 0.0001). Octoploid samples exhibited a higher proportion of unmethylated sites compared to tetraploid samples (0.12% vs. 0.08%), while the proportion of fully methylated sites was lower in octoploids than in tetraploids (0.37% vs. 0.41%). PCoA revealed substantial, though incomplete, separation of samples by ploidy level (Fig. 4b). Genotypes Ru8x and Hu8x had the most distinct methylation patterns (Suppl. Fig. 1).

## 4. Discussion

### 4.1. Response to drought

The molecular response to drought in plants involves a complex network of pathways and gene expression changes, which interact with other stress response mechanisms, creating a highly integrated system. Several key proteins have been previously identified in drought-stressed plants, including late embryogenesis abundant (LEA) proteins, desiccation stress proteins, abscisic acid-responsive proteins, enzymes involved in the synthesis of osmoprotectants (sugars, proteins, glycine betaine), cold-regulated proteins, detoxification enzymes, as well as numerous transcription factors (Tzvi *et al*. 2000).

We identified multiple up-regulated genes previously reported to be associated with drought stress. Among them, two showed the highest up-regulation on both the 20th and 30th days of drought across both ploidy levels (Fig. 3a). The first gene encodes alpha-aminoadipic semialdehyde synthase (AASS, upregulated 5.0- to 8.4-fold), an enzyme that plays a crucial role in lysine catabolism through the saccharopine pathway in plants. A gene of another protein from this pathway, saccharopine dehydrogenase (*SCCPDH,* IPR032095), was also up-regulated for these groups of samples. The saccharopine pathway has been previously shown to be upregulated under osmotic stress, contributing to the production of the osmolytes proline and pipecolate, which aid in water stress adaptation (Rodrigues *et al*. 2006; Arruda and Barreto 2020). Although AASS is not typically classified among the key water-stress proteins, its gene has been identified among a hub of up-regulated genes under water stress in *Nicotiana tabacum* and *Hordeum vulgare* (Chen *et al*. 2019; Panahi and Golkari 2024).

The second most upregulated transcript (4.7- to 8.0-fold) encodes the LEA1 protein (IPR005513). LEA proteins are classified into eight subgroups of small, highly hydrophilic polypeptides: LEA1 – LEA6, dehydrin, and seed maturation protein (SMP) (Hunault and Jaspard 2010). LEA proteins play a crucial role in protecting plant metabolism under abiotic stresses such as drought. Progressive drought and salinity stress induce LEA proteins in vegetative organs, where they contribute to stabilizing enzyme complexes and maintaining cell membrane integrity through antioxidant activity, metal ion binding, membrane and protein stabilization, hydration buffering, and interactions with DNA and RNA (Chourey *et al*. 2003; Goyal *et al*. 2005; Battaglia and Covarrubias 2013; Liang *et al*. 2019). Additionally, two genes encoding other LEA proteins - dehydrin (IPR000167) and SMP (IPR007011) - were also found to be upregulated up to 4.4-fold and 3.0-fold, respectively.

A third most highly upregulated (up to 6.6-fold) gene under drought stress is encoding endo-1,4-β-xylanase (IPR001000), an enzyme involved in remodelling the plant cell wall by hydrolysing xylan. Knockout studies in *Zea mays* have shown that loss of this gene results in thinner cell walls with reduced xylan and lignin content compared to wild-type plants. These structural deficiencies ultimately impaired water transport capacity and lead to lower drought tolerance (Hu *et al*. 2020).

One more pathway was found to be involved in the drought response. Three different up-regulated transcripts were assigned to the gene *HMGCR* encoding hydroxymethylglutaryl-CoA reductase (IPR009029, IPR009023, IPR002202) a key enzyme in the mevalonate pathway for isoprenoid biosynthesis. Isoprenoids play an important role in plant stress responses by providing protection against oxidative damage, thermal stress, and drought, while also contributing to stress signalling and gene regulation (Vickers *et al*. 2009; Zuo *et al*. 2019). Interestingly, *HMGCR* expression was significantly higher on the 20th day of drought compared to the 30th day, suggesting dynamic changes in response to increasing drought severity. This pattern aligns with the observations of Perreca *et al*. (2020), who reported that isoprene content in *Picea glauca* needles increased under mild drought stress but decreased under more severe drought conditions.

The pathways involved in drought and salt stress responses are known to substantially overlap in plants (Ha *et al*. 2014; Krasensky and Jonak 2012; Luo *et al*. 2015). Therefore, we expected gene expression patterns similar to those reported in previous studies on *P. australis* under salinity stress (Holmes *et al*. 2016; Jia *et al*. 2024). Surprisingly, only one gene, *dehydrin*, was consistently identified among the core up-regulated genes. However, consistent with these studies and a study on *P. australis* grown under low water levels (Eller *et al*. 2014; Haldan *et al*. 2023), no oxidative stress response genes, such as manganese superoxide dismutase (IPR019832/IPR019831/IPR036324) or glutathione peroxidase (IPR000889), were up-regulated.

We identified additional up-regulated genes that are potentially involved in drought stress. These genes encode non-symbiotic haemoglobin (*NsHb*, IPR000971), oil body-associated protein-like (*OBAP-like*, IPR010686), Bowman-Birk inhibitor (*BBI*, IPR000877), osmotin (IPR037176), heat shock associated protein 32 (*HSA32*, IPR003830), β-fructofuranosidase (IPR021792), novel interactor of jasmonoyl (*NINJA*) or ABI five-binding protein (*AFP*) (both IPR032310), cold-regulated protein 413 (*COR413*, IPR008892), glycoside hydrolase (GH3, IPR002772), glyoxalase (*GLO*, IPR029068), and tyrosinase (Cu-binding domen, IPR002227).

### 4.2. Photosynthesis under drought

Drought is known to suppress photosynthetic activity in plants by disrupting the balance between light capture and its utilization (Foyer and Noctor 2000). Previous studies have shown that *P. australis* tolerates mild water deficits by reducing leaf area while maintaining photosynthetic efficiency and Rubisco activity through increased water use efficiency (Pagter *et al*. 2005; Nada *et al*. 2015; Liu *et al*. 2018). However, more severe drought conditions impose biochemical limitations (Teng *et al*. 2022). In our previous mesocosm experiment, *P. australis* plants were grown with the water level 45 cm below the soil surface and exhibited no decrease in photosynthesis rate and expression of three key photosynthetic genes (*Ribulose bisphosphate carboxylase, Phosphoglycerate kinase,* and *Phosphoribulokinase*) compared to those with water at the soil level (Haldan *et al*. 2023). That could suggest that the decrease in water availability was not severe enough to induce stress.

In the current study, we observed a progressive decline in the photosynthetic rate throughout the drought period, with the most pronounced decrease occurring within the first 20 days (Fig. 3c in Haldan *et al*. 2025). We observed that one of the genes, *PsbP*, was substantially down-regulated, with a 22.7- and 34.9-fold change after 20 and 30 days of drought, respectively (Fig. 3b). This gene encodes the P subunit of the oxygen-evolving complex in Photosystem II (PSII), which oxidizes water to provide protons for Photosystem I (PSI). PsbP subunit is thought to play a role in regulating the balance between water oxidation and plastoquinone reduction (Ifuku and Nagao 2021). Plastoquinone acts as an antioxidant, and its reduction status has been recognized as a key indicator of stress impact on photosynthesis (Havaux 2020, Sperdouli *et al*. 2021).

More photosynthetic genes were found to be down-regulated, though less strong. After 20 days of drought, *PsaO*, *PsaF*, and *carbonic anhydrase* were down regulated (2.3-, 2.3-, 4.3-, and 1.7-fold, respectively). On the 30th day of drought, *PsbO* (only for tetraploid samples) and *crr42* (protein chlororespiratory reduction) were down-regulated 4.3- and 3.2-fold, respectively. Gene ontology enrichment analysis revealed that on the 20th day of drought, the cellular component terms PSI and PSI reaction center were among the down-regulated transcripts. On the 30th day of drought, terms related to the assembly of the NADH dehydrogenase complex (plastoquinone) and the mitochondrial respiratory chain complex I were identified.

### 4.3. DNA methylation pattern under drought

It was previously shown that plants can exhibit increased variance in DNA methylation when exposed to ecological stressors (Ashapkin *et al*. 2020; Akhter *et al*. 2021). For example, genome-wide hypermethylation occurs in drought-stressed plants like *Morus alba* and *Populus trichocarpa*, with differentially methylated regions linked to stress-response genes (Sun *et al*. 2022). Moreover, it was shown that drought-tolerant varieties of these species maintain more stable methylomes, while drought-sensitive plants show greater methylation variability.

Despite our AMOVA results showed no significant differences between drought treatments, the PCoA revealed an increase in variance among samples as drought progressed. By the 40th and 50th days of drought, methylation patterns were the most scattered and the least scattered in control samples, highlighting genotype-specific differences in drought response.

Previous epigenetic studies on *P. australis* (but not using MSAP) showed that DNA methylation is implicated in response to salinity. Results indicated that DNA methylation levels tend to decrease with increased salinity (Petroff 2013; Song *et al*. 2022). We found no statistically significant correlation between drought duration and methylation level, however, similar to previous studies, the pattern was positive for unmethylation (r = 0.67) and negative for full methylation (r = -0.64) levels.

### 4.4. Difference between ploidy levels

Polyploid plants are often, though not always, shown to exhibit superior drought tolerance compared to their diploid counterparts, attributed to enhanced abscisic acid signalling, improved photosynthetic and hormonal regulation, and increased antioxidant activity (Allario *et al*. 2013; Xue *et al*. 2015; Rao *et al*. 2020; Tossi *et al*. 2022; Correia *et al*. 2023). Their resilience under extreme stress conditions may also be linked to flexible reproductive strategies and the production of heavier, more viable seeds, particularly in grasses (Godfree *et al*. 2017; Stevens *et al*. 2020). However, our data on growth and photosynthesis, along with other common garden experiments on *P. australis* (Hansen *et al*. 2007; Achenbach *et al*. 2012), provide no evidence that octoploid plants outperform tetraploid plants under the same or stressful environmental conditions.

Previous transcriptomic analysis of three tetraploid and three octoploid genotypes identified differences in gene expression: tetraploid genotypes up-regulated genes related to reproduction and defence against UV-B light and fungi, while octoploid genotypes up-regulated genes associated with thermotolerance (Wang *et al*. 2021). However, no regional pairs were included in this study — all three octoploid plants originated from Australia, while tetraploid plants were collected from North America and Europe. Given the complex phylogenetic relationships within the species and the high variation in phenotypic plasticity and gene expression among genotypes and lineages (Takahashi *et al*. 2007; Achenbach *et al*. 2012; Eller *et al*. 2014; Zhang *et al*. 2020; Haldan *et al*. 2023), the observed differences may be driven by genotype-specific variation and local adaptation. To ensure robust comparisons of *P. australis* in future studies, a broader range of genotypes from diverse origins should be included.

Higher ploidy can significantly impact transcription and translation regulation networks due to increased genomic complexity. In our study, up-regulated genes identified in octoploid samples are mostly involved in translation regulation, but also in protein binding, and biosynthesis. These enriched biological processes can mitigate proteotoxic stress, ensure fidelity in protein synthesis and support metabolic demands of larger cells. The most highly upregulated gene (7.2-fold) encodes the Mago Nashi protein, which plays a role in RNA metabolism, developmental signalling and stress response in plants (He *et al*. 2007). To date, there is no direct evidence on ploidy-associated expression of this gene in literature, but empirical studies are needed. Two genes found up-regulated in tetraploid samples are involved in cell wall modification and transmembrane transport.

Tetraploid and octoploid samples responded differently to the drought. Between the 20th and 30th days of drought, tetraploid samples reduced the number of down-regulated transcripts, while octoploid samples reduced the number of up-regulated transcripts (Fig. 2a).

AMOVA and PCoA of MSAP data revealed differences in DNA methylation between tetraploid and octoploid samples, with octoploids exhibiting lower methylation levels. This difference is likely driven by genotype- or ploidy-related factors rather than drought response, as variation between treatments was smaller than that between genotypes (Suppl. Fig. 1). Another study using the MSAP method found that triploid *Citrullus vulgaris* and *Salvia miltiorrhiza* had 18 – 22% lower methylation levels than their diploid counterparts, whereas triploid *Populus* and *Pyrus × bretschneideri* exhibited 24 – 25% higher methylation levels than their diploid parents (Li *et al*. 2011). The absence of a linear relationship between ploidy and methylation levels suggests a complex, species-specific epigenetic regulation. Additionally, evolutionary timing and hybridization type (allo-vs. autopolyploids) have a strong influence on methylation patterns (Yaakov and Kashkush 2011; Duan *et al*. 2023). Focusing on epigenetic modifications of stress-response genes, rather than genome-wide methylation, may provide more precise insights into the role of epigenetic changes in the stress adaptation of *P. australis* as a first step toward developing epigenetic control strategies for future hybridization and selection studies.

## Conclusions

The molecular response to drought in *P. australis* involves a complex network of gene expression changes. We identified several key drought-response genes common to both ploidy levels, including those involved in saccharopine pathway (*AASS* and *SCCPDH*), response to water deprivation (*dehydrin* and *LEA1*), cell wall remodelling (*endo-1,4-β-xylanase*), and mevalonate pathway (*HMGCR*). Drought suppressed photosynthesis, with a drastic down-regulation of *PsbP*. Polyploidy influenced gene expression patterns, with octoploid samples up-regulating genes involved in translation and metabolism, while tetraploids activated genes associated with cell wall modification and transmembrane transport. Ploidy levels displayed distinct gene expression dynamics throughout drought stress. Under non-stress conditions, a distinct gene expression pattern was observed, with different genes being differentially regulated. Notably, octoploid plants exhibited the strongest upregulation of the gene encoding the Mago Nashi protein. DNA methylation patterns showed increased variability under prolonged drought, although no significant correlation was found between methylation levels and drought duration. Pronounced differences in DNA methylation were found between tetraploid and octoploid plants, with octoploids exhibiting a lower methylation level. Given the complex phylogenetic relationships and genotype-specific responses of *P. australis*, future studies should incorporate a broader range of genotypes to improve comparisons. Additionally, focusing on epigenetic modifications and other regulation mechanisms of stress-response genes, could provide new insights into drought adaptation and potential epigenetic control strategies for hybridization, selection and conservation studies.

## Acknowledgements

Many thanks to Dr. Franziska Eller from Aarhus University for kindly providing information about the samples, and to Robin Landeau and Anja Klahr for their assistance in establishing the MSAP method. We are grateful to Kai-Uwe Schwarz and Heike Meyer from the Julius-Kühn Institute, Braunschweig, for the meristematic propagation of plants.

## Author Contribution

**Kristina Kuprina:** Conceptualization, Investigation, Data curation, Statistical analysis, Visualization, Writing - original draft. **Kerstin Haldan**: Conceptualization, Investigation, Writing - review and editing. **Stepan Saenko:** Investigation, Statistical analysis, Writing - review and editing. **Gulam Mohamed Safwaan:** Investigation, Writing - review and editing. **Jürgen Kreyling:** Conceptualization, Writing - review and editing. **Martin Schnittler:** Conceptualization, Funding acquisition, Supervision, Writing - review and editing: **Manuela Bog:** Conceptualization, Supervision, Writing - review and editing.

## Data availability

Clean short and long reads, as well as assembled transcriptome, can be found on NCBI (PRJNA1195549). The details on data processing and analysis can be found on GitHub: https://github.com/kuprinak/Phragmites_drought_RNAseq.

