## Supplementary material for "How drought and ploidy level shape gene expression and DNA methylation in *Phragmites australis*": Suppl.Fig.1

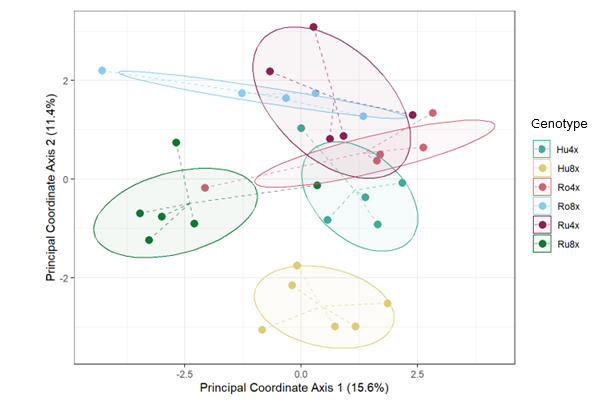

Supplementary Figure 1. Ordination diagram of PCoA calculated on 248 methylation-sensitive loci for 32 leaf samples of six genotypes of *Phragmites australis*. Ellipses depict one standard deviation of the data (α = 0.68).
